# Coagulation factors directly cleave SARS-CoV-2 spike and enhance viral entry

**DOI:** 10.1101/2021.03.31.437960

**Authors:** Edward R. Kastenhuber, Javier A. Jaimes, Jared L. Johnson, Marisa Mercadante, Frauke Muecksch, Yiska Weisblum, Yaron Bram, Robert E. Schwartz, Gary R. Whittaker, Lewis C. Cantley

## Abstract

Coagulopathy is a significant aspect of morbidity in COVID-19 patients. The clotting cascade is propagated by a series of proteases, including factor Xa and thrombin. While certain host proteases, including TMPRSS2 and furin, are known to be important for cleavage activation of SARS-CoV-2 spike to promote viral entry in the respiratory tract, other proteases may also contribute. Using biochemical and cell-based assays, we demonstrate that factor Xa and thrombin can also directly cleave SARS-CoV-2 spike, enhancing viral entry. A drug-repurposing screen identified a subset of protease inhibitors that promiscuously inhibited spike cleavage by both transmembrane serine proteases as well as coagulation factors. The mechanism of the protease inhibitors nafamostat and camostat may extend beyond inhibition of TMPRSS2 to coagulation-induced spike cleavage. Anticoagulation is critical in the management of COVID-19, and early intervention could provide collateral benefit by suppressing SARS-CoV-2 viral entry. We propose a model of positive feedback whereby infection-induced hypercoagulation exacerbates SARS-CoV-2 infectivity.

## Introduction

SARS-CoV-2 emerged into the human population in late 2019 and has evolved into a devastating global health crisis. Despite the recent success of vaccines (Baden et al., 2020; Polack et al., 2020), the limited world-wide vaccine distribution (Kwok et al., 2020; Lin, Tu, & Beitsch, 2020; Nhamo, Chikodzi, Kunene, & Mashula, 2020; So & Woo, 2020), the emergence of viral variants (Wang et al., 2021; Weisblum et al., 2020), and the repeated SARS-like zoonotic outbreaks over the last 20 years (Cheng, Lau, Woo, & Yuen, 2007; Ge et al., 2013; Menachery et al., 2015) underscore the urgent need to develop antivirals for coronavirus (Consortium et al., 2020).

In addition to attachment to specific receptors on target cells, coronaviruses require proteolytic processing of the spike protein by host cell proteases to facilitate membrane fusion and viral entry (Glowacka et al., 2011; J. A. Jaimes, J. K. Millet, & G. R. Whittaker, 2020; Walls et al., 2020). In SARS-CoV-2, host cell proteases act on two sites residing at the S1/S2 subunit boundary and at the S2’ region proximal to the fusion peptide (Belouzard, Chu, & Whittaker, 2009; Hoffmann, Kleine-Weber, et al., 2020; J. Jaimes, J. Millet, & G. Whittaker, 2020; Millet & Whittaker, 2014). S1/S2 site cleavage opens up the spike trimer and exposes the S2’ site, which must be cleaved to allow for the release of the conserved fusion peptide (Benton et al., 2020; Madu, Roth, Belouzard, & Whittaker, 2009). While the prevailing model suggests that furin cleaves the S1/S2 site and TMPRSS2 cleaves the S2’ site (Bestle et al., 2020), it remains unclear to what extent other proteases may be involved (Hoffmann et al., 2021; Ou et al., 2020).

TMPRSS2 is an important host cell factor in proteolytic activation across multiple coronaviruses (Hoffmann, Kleine-Weber, et al., 2020; Jaimes, Millet, Goldstein, Whittaker, & Straus, 2019). TMPRSS2 knockout or inhibition reduces infection in mouse models of SARS and MERS (Iwata-Yoshikawa et al., 2019; Y. Zhou et al., 2015). More recently, TMPRSS2 has been highlighted as a drug target for SARS-CoV-2 (Hoffmann, Kleine-Weber, et al., 2020; Hoffmann, Schroeder, et al., 2020).

Furin activity is not essential to produce infectious particles (Tang et al., 2021) and furin is not necessary for cell fusion (Papa et al., 2021), but deletion of the S1/S2 site attenuates SARS-CoV-2 *in vivo* (Johnson et al., 2021). Proteolytic activation of envelope proteins presumably coordinates target cell engagement and envelope conformational changes leading to fusion. Furin cleavage during viral biogenesis, before release of viral particles, may render SARS-CoV-2 spike less stable in solution and reduce the likelihood to reach and interact with target cells (Amanat et al., 2021; Berger & Schaffitzel, 2020; Wrobel et al., 2020). Although the S1/S2 site is often referred to as the “furin site” (Johnson et al., 2021), the full spectrum of proteases that catalyze biologically relevant activity in the lung remains incompletely defined.

Proteases also orchestrate the coagulation pathway, via a series of zymogens that are each activated by a chain reaction of proteolytic processing. Coagulopathy and thromboembolic events have emerged as a key component of COVID-19 pathogenesis (McGonagle, O’Donnell, Sharif, Emery, & Bridgewood, 2020). Comorbidities associated with severe COVID-19 are also linked to dysregulated blood clotting (F. Zhou et al., 2020). Patients with a history of stroke prior to infection have nearly twice the risk of in-hospital mortality (Qin et al., 2020). Upon hospital admission, elevated D-dimer levels (an indicator of fibrinolysis and coagulopathy) and low platelet counts (an indicator of consumptive coagulopathy) are predictive biomarkers of severe disease and lethality in COVID-19 patients (F. Zhou et al., 2020). Systemic activity of clotting factors V, VIII, and X are elevated in severe COVID-19 disease (Stefely et al., 2020). While early phase disease is typically restricted to a local pulmonary hypercoagulable state, late stage disease may be accompanied by systemic DIC, stroke and cardio-embolism (Huang et al., 2020; Kipshidze et al., 2020; McGonagle et al., 2020; Tsivgoulis et al., 2020). Ischemic stroke occurred in approximately 1% of hospitalized COVID-19 patients, and strikingly, a fraction of them experienced stroke even prior to onset of respiratory symptoms (Yaghi et al., 2020).

In a drug repurposing effort to target TMPRSS2, we observed that multiple direct-acting anticoagulants have anti-TMPRSS2 off-target effects. We subsequently investigated overlap in substrate specificity between TMPRSS2, factor Xa and thrombin. Circulating proteases involved in blood clotting can cleave and activate SARS-CoV-2 spike, enhancing viral entry. We propose that the serine protease inhibitor nafamostat may incorporate a combined mechanism in the treatment of COVID-19 through inhibition of TMPRSS2 and coagulation factors.

## Results

### Serine protease inhibitors suppress SARS-CoV-2 entry via inhibition of TMPRSS2

We developed a fluorescence resonance energy transfer (FRET)-based protease enzymatic assay based on peptides containing either the S1/S2 or S2’ cleavage sites of SARS-CoV-2 spike (**Fig. 1A, S1**). Upon cleavage, the liberated 5-FAM emits fluorescent signal proportional to the quantity of product (**Fig. S1A**). Camostat and nafamostat resulted in strong inhibition of TMPRSS2 (**Fig. 1B**), as expected (Hoffmann, Kleine-Weber, et al., 2020; Hoffmann, Schroeder, et al., 2020). We also identified that otamixaban and the active form of dabigatran (but not its prodrug dabigatran etexilate) inhibit TMPRSS2 enzymatic activity *in vitro* (**Fig. 1B-C**).

**Figure 1.**
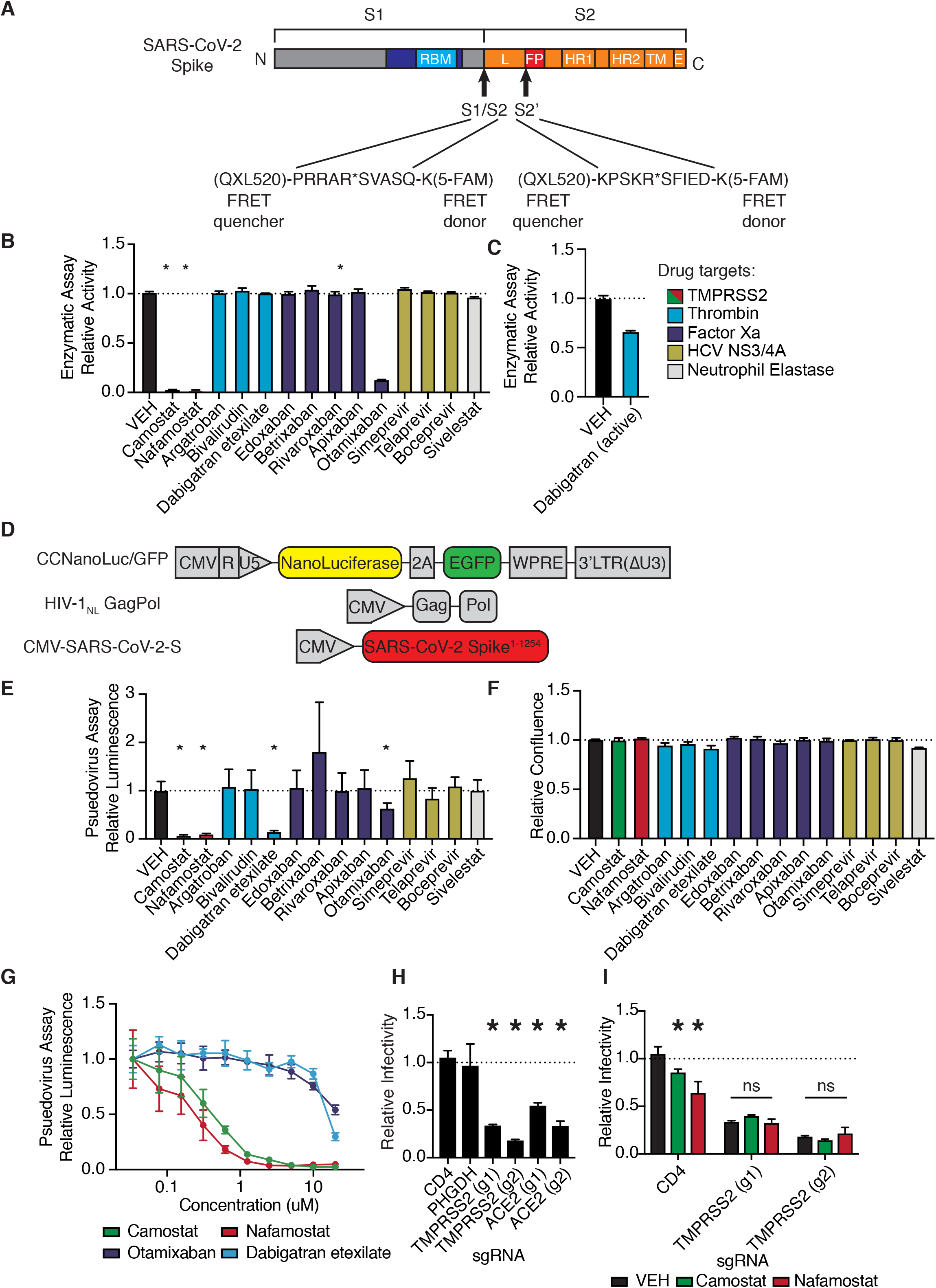
Anticoagulant serine protease inhibitors suppress SARS-CoV-2 entry via inhibition of TMPRSS2. (**A**) Peptides derived from two known cleavage sites of SARS-CoV-2 spike were designed with C-terminal fluorophore 5-FAM and N-terminal FRET quencher QXL-520. (**B**) FDA-approved and investigational serine protease inhibitors were screened by enzymatic assay to inhibit TMPRSS2 cleavage of SARS-CoV-2 S1/S2 peptide substrate. Relative change in fluorescence with respect to DMSO vehicle is shown. Colors indicate the described target of the drugs screened. All drugs screened at 10 *µ*M final concentration. (**C**) Active form of dabigatran in enzymatic assay for TMPRSS2 inhibition. Relative fluorescence with respect to its corresponding 0.1N HCl vehicle is shown. (**D**) Schematic of constructs used to generate SARS-CoV-2 spike-pseudotyped/HIV-1-based particles. (**E**) Calu3 cells were treated with 10 *µ*M of the indicated drugs for 24 hr prior to infection with HIV-1_NL_/SARS-CoV-2 pseudovirus. Media was changed at 24 hr post infection and pseudoviral entry was measured by Nanoluciferase luminescent signal at 40 hr. (**F**) Calu3 cells treated with 10 *µ*M of the indicated drugs were monitored for confluence by Incucyte for 40 hr. (**G**) Pseudoviral entry was measured by Nanoluciferase luminescent signal in Calu3 cells treated various concentrations of the indicated drugs for 4 hours prior to infection with SARS-CoV-2 pseudovirus. (**H**) Caco2 cells were infected with lenti-Cas9-blast and U6-sgRNA-EFS-puro-P2A-tRFP and selected. Neutral controls targeting CD4 (not endogenously expressed) or PHGDH intron 1, two sgRNAs each targeting different regions of ACE2 and TMPRSS2 were included. Cells were subsequently infected with HIV-1_NL_/SARS-CoV-2 pseudovirus. (**I**) Caco2 cells co-expressing Cas9 and sgRNAs targeting CD4 (not expressed) or TMPRSS2 were treated with 10*µ*M Camostat, Nafamostat, or DMSO vehicle. * P<0.05, two-tailed t-test. Error bars +/− SEM.

To explore these candidates in a cell-based functional assay of spike protein, SARS-CoV-2 S-pseudotyped HIV-1 particles were employed to infect human lung Calu3 cells (**Fig. 1D**) (Schmidt et al., 2020). Consistent with the TMPRSS2 enzymatic assay, camostat, nafamaostat, otamixaban, and dabigatran etexilate suppressed pseudoviral entry, as indicated by nanoluciferase luminescent signal (**Fig. 1E**). No effects on relative cell growth were observed at the same timepoint in Calu3 (**Fig. 1F)** or A549 cells (data not shown), confirming that reduced luminescent signal was not due to cytotoxicity. A dose-response experiment with select protease inhibitors revealed a sub-micromolar IC50 for camostat and nafamostat and IC50s in the 10-20 *µ*M range for otamixaban and dabigatran in Calu3 cells (**Fig. 1G**).

Using A549 cells with or without ectopic ACE2 expression, we confirmed that HIV-1_NL_/SARS-CoV-2 pseudovirus infection is dependent on ACE2, while infection with HIV-1_NL_ pseudotyped instead with VSV G envelope protein is not ACE2 dependent (**Fig. S2**). Caco2 cells, which endogenously express ACE2 and TMPRSS2, show greater susceptibility to SARS-CoV-2 S-pseudotyped HIV-1_NL_, but equivalent susceptibility to VSV G-pseudotyped HIV-1_NL_, when compared to A549/ACE2 cells (**Fig. S2**).

To further validate these results in an alternative pseudovirus system, we used recombinant G protein-deficient vesicular stomatitis virus (rVSVΔG) pseudotyped with SARS-CoV-2-S (**Fig. S3A**), yielding pseudovirus dependent on spike for cell entry (**Fig. S3B**). The antiviral effects of the four candidate protease inhibitors were confirmed in the VSV pseudovirus system in multiple cell lines, and response was associated with TMPRSS2 expression (**Fig. S3C-F**).

We aimed to determine whether the effects of camostat and nafamostat are indeed TMPRSS2-dependent, or if other unidentified cellular proteases can compensate for TMPRSS2 suppression. To do so, we knocked out TMPRSS2 in ACE2^+^TMPRSS2^+^ Caco2 cells and found that susceptibility to pseudovirus was significantly reduced, comparable to knockout of ACE2 (**Fig. 1H, S4**). Furthermore, both camostat and nafamostat reduce pseudovirus entry into control Caco2 cells harboring control sgRNA, but this effect was abolished in cells with two independent TMPRSS2-targeting sgRNAs (**Fig. 1I**). These data indicate that, in the absence of exogenous proteases, TMPRSS2 is a critical host enzyme activating SARS-CoV-2 spike in TMPRSS2^+^ cells and that TMPRSS2 is the primary target of camostat and nafamostat in these conditions.

### Coagulation factors directly cleave SARS-CoV-2 spike

Anticoagulants are highly represented among FDA-approved drugs that target proteases, and among the hits from the screen described above. The off-target effects of anticoagulants on TMPRSS2 imply that these small molecules can interact with the active sites of TMPRSS2 in a similar manner to coagulation factors. This led us to hypothesize that coagulation factors may interact with some of the same substrates as TMPRSS2, including SARS-CoV-2 spike.

To determine the properties of enzyme-substrate relationships, TMPRSS2, factor Xa, and thrombin cleavage of S1/S2 and S2’ peptides were determined over a range of 0-160 *µ*M peptide substrate (**Fig. 2A-C, Table 1**). Surprisingly, factor Xa catalyzed S1/S2 cleavage more than an order of magnitude faster than TMPRSS2 (**Fig. 2A-B,D**), although factor Xa showed lower affinity (higher K_m_) compared with TMPRSS2 to the S1/S2 peptide (**Fig. 2A-B,E**). Thrombin has greater affinity (lower K_m_) than TMPRSS2 and Factor Xa for the S1/S2 substrate (**Fig. 2E**) and performs S1/S2 cleavage at a rate intermediate between TMPRSS2 and Factor Xa (**Fig. 2D**). Unlike factor Xa, thrombin cleaves the S2’ peptide with greater activity than TMPRSS2 (**Fig. 2D-F**).

**Figure 2.**
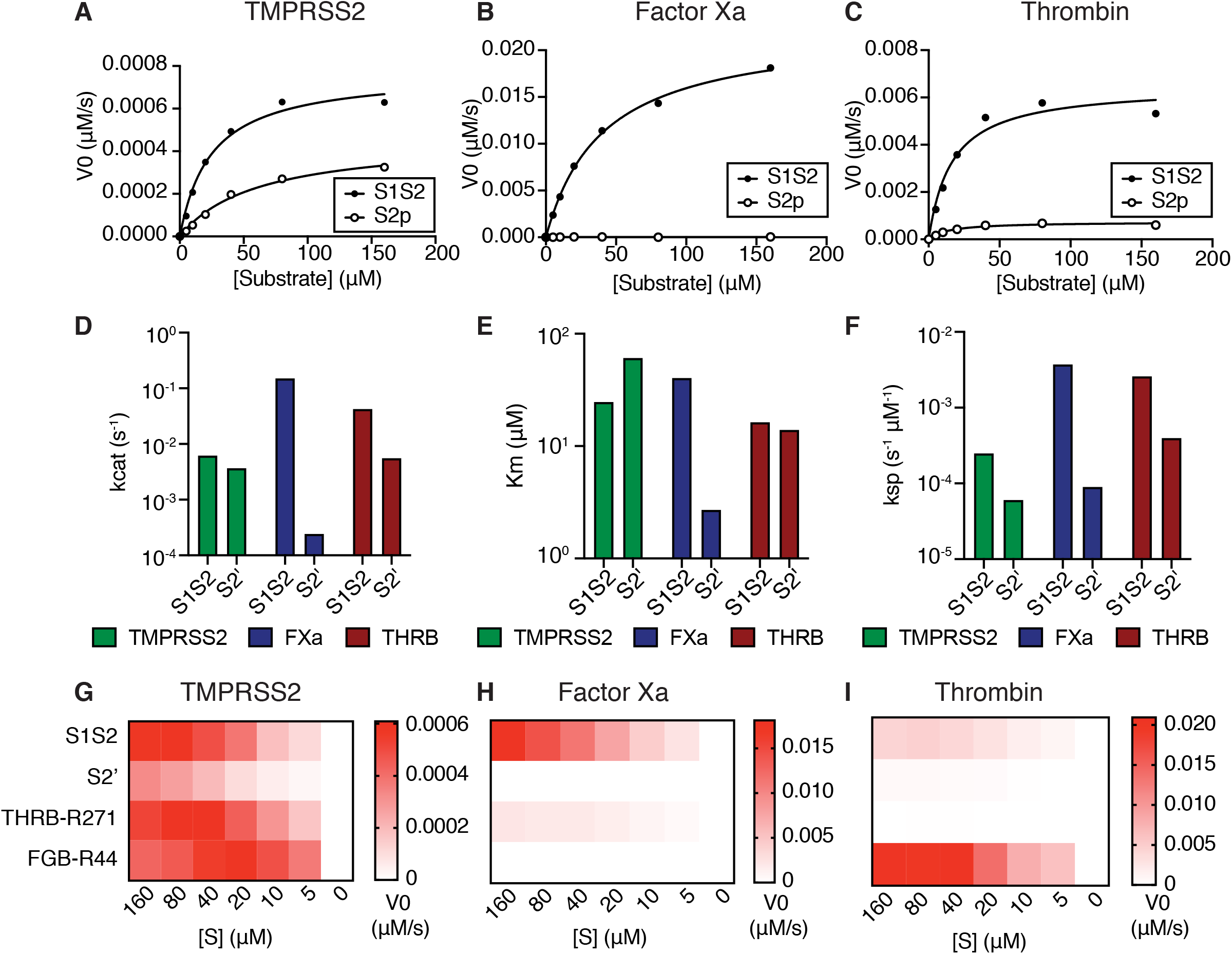
Coagulation factors directly cleave SARS-CoV-2 spike. Initial velocities for the cleavage of SARS-CoV-2 spike S1/S2 and S2’ peptide substrates by (**A**) TMPRSS2, (**B**) Factor Xa, and (**C**) Thrombin were measured over a range of 0-160 *µ*M substrate. From initial velocity values, enzyme kinetic constants (**D**) turnover rate K_cat_ (s^−1^), (**E**) affinity constant K_m_, and (**F**) specificity constant (K_cat_/K_m_) were obtained for the indicated enzymes with S1/S2 and S2’ peptides. (**G-I**) Heatmaps depict the initial velocity V_0_ of cleavage of the indicated peptide substrates and concentrations by (**G**) TMPRSS2, (**H**) factor Xa, and (**I**) thrombin.

**Table 1.** Kinetics of SARS-CoV-2 spike peptide substrate cleavage. Kinetic constants obtained from initial velocity studies with varying concentrations of SARS-CoV-2 spike S1/S2 and S2’ peptide substrates. Each estimate is based on seven different concentrations of substrate in 1:2 serial dilution (0-160 *µ*M).

| **Enzyme** | **Substrate** | **$V_{\max}$ ($\mu\text{M/s}$)** | **$k_{\text{cat}}$ ($\text{s}^{-1}$)** | **$K_{\text{m}}$ ($\mu\text{M}$)** | **$K_{\text{sp}}$ ($\text{s}^{-1} \mu\text{M}^{-1}$)** |
| --- | --- | --- | --- | --- | --- |
| TMPRSS2 | S1/S2 | 7.71E-04 | 6.17E-03 | 24.71 | 2.50E-04 |
| TMPRSS2 | S2' | 4.60E-04 | 3.68E-03 | 60.94 | 6.04E-05 |
| Factor Xa | S1/S2 | 2.24E-02 | 1.79E-01 | 40.35 | 4.43E-03 |
| Factor Xa | S2' | 3.04E-05 | 2.43E-04 | 2.711 | 8.97E-05 |
| Thrombin | S1/S2 | 6.50E-03 | 5.20E-02 | 16.34 | 3.18E-03 |
| Thrombin | S2' | 7.34E-04 | 5.87E-03 | 13.98 | 4.20E-04 |

We next compared the ability of coagulation factors to cleave SARS-CoV-2 S to their ability to cleave their known substrates. During the physiological process of clotting, factor Xa cleaves prothrombin at R271, which ultimately becomes the activated form *α*-thrombin (Wood, Silveira, Maille, Haynes, & Tracy, 2011). Thrombin, in turn, cleaves multiple sites of fibrinogen, including the beta chain (FGB) at R44, in a critical step toward aggregation and polymerization of high molecular weight fibrin clots. Fluorogenic peptides corresponding to THRB^R271^ and FGB^R44^ were synthesized and assayed with TMPRSS2, factor Xa, and thrombin. TMPRSS2 exhibited relatively broad activity to cleave this collection of substrates (**Fig. 2G**). As expected, factor Xa showed strong selectivity for THRB^R271^ over FGB^R44^, while thrombin showed the opposite substrate preference (**Fig. 2H-I**). Remarkably, factor Xa showed ∼9-fold greater maximum initial reaction velocity (V_max_) in cleaving the spike S1/S2 peptide compared to cleaving its known substrate, THRB^R271^ (**Fig. 2H**). The V_max_ for thrombin cleavage of the spike S1/S2 peptide was only ∼4.5-fold lower than its V_max_ for the benchmark FGB^R44^ peptide (**Fig. 2I**), indicating that thrombin might also cleave this site when activated during coagulation.

We next assessed the effect of substituting amino acids adjacent to the cleavage site of the S1/S2 peptide on proteolytic cleavage by these proteases. An arginine preceding the cleavage site (P1 position) is a common feature of substrates of many serine proteases. Substitution of the P1 arginine in the S1/S2 substrate with alanine (S1S2-P1A) resulted in a 4-fold reduction in TMPRSS2 cleavage and abolished nearly all cleavage by factor Xa and thrombin (**Fig. S5A-C**). Substitutions in the P3 and P4 positions (S1S2-HPN) with features typical of a substrate of type II transmembrane serine proteases (TTSPs), a family which includes TMPRSS2 and hepsin (Damalanka et al., 2019) did not change TMPRSS2 cleavage and greatly reduced factor Xa and thrombin cleavage (**Fig. S5A-C**). Although the substrate specificity of TTSPs and coagulation factors are not uniformly similar, the sequence of the SARS-CoV-2 S1/S2 site is a substrate common to TMPRSS2, factor Xa, and thrombin.

In summary, the coagulation serine proteases factor Xa and thrombin exhibit even greater proteolytic activity against the SARS-CoV-2 peptide substrates than TMPRSS2, a protease essential in coronavirus entry, and the S1/S2 boundary appears to be an even more optimal factor Xa substrate than peptide substrates derived from known physiological targets of factor Xa in coagulation.

### Factor Xa and thrombin facilitate SARS-CoV-2 spike mediated entry

We next investigated whether coagulation factors could cleave trimeric spike in its native 3D conformation, and whether this activity potentiated spike function in viral entry into cells. To do so, we used replication-defective SARS-CoV-2 spike-pseudotyped VSV or HIV-1 virus (Schmidt et al., 2020). In the VSV pseudovirus system, addition of purified factor Xa or thrombin to the media significantly increased infection in Calu3 cells 16 hours post infection as determined by either quantification of NeonGreen (**Fig. 3A, S6A**) or Nanoluciferase activity **(Fig. 3B)**.

**Figure 3.**
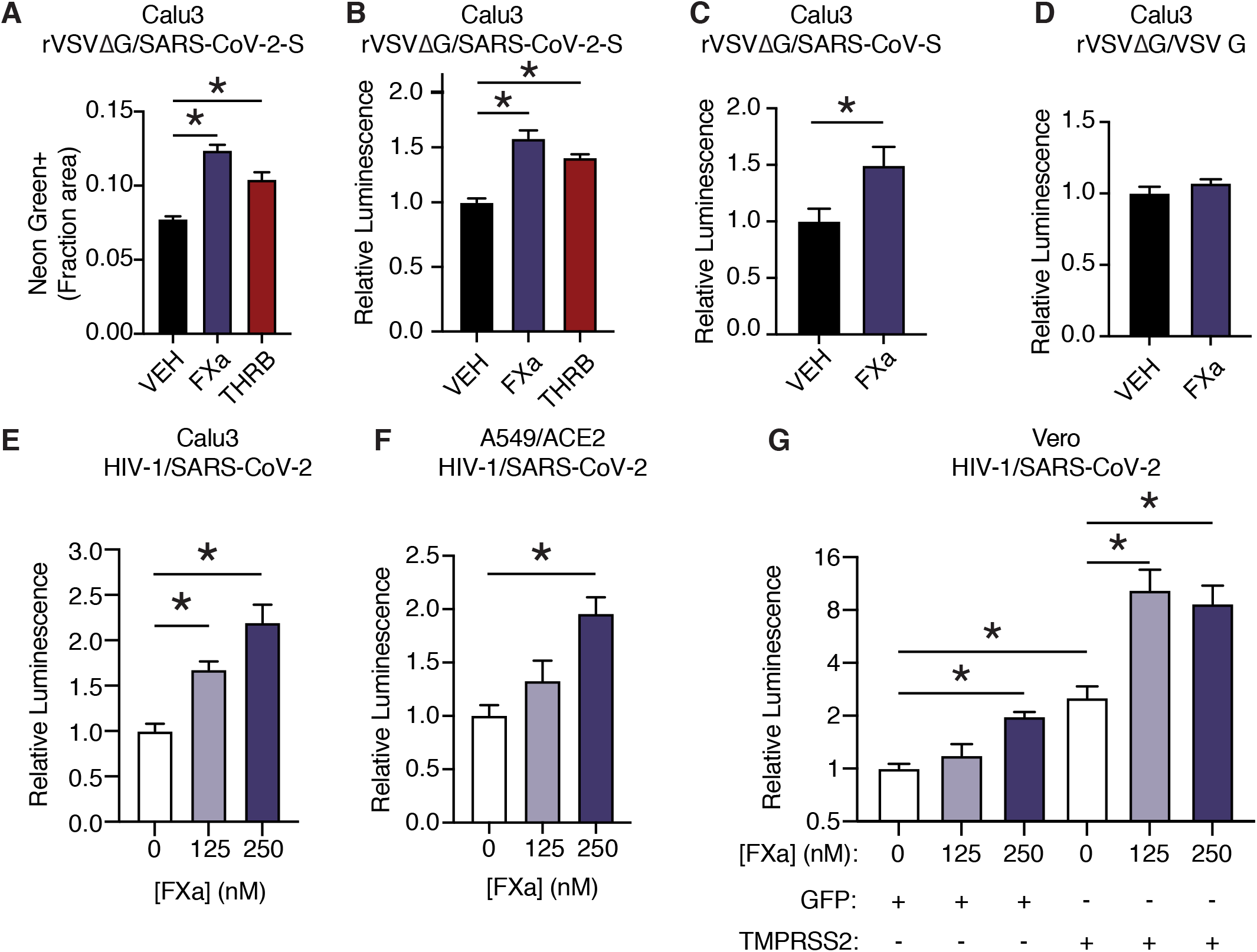
Factor Xa and thrombin facilitate SARS-CoV-2 spike mediated entry. (**A**) Calu3 cells were infected with rVSVΔG/SARS-CoV-2 pseudovirus with concomitant treatment with vehicle, 250nM factor Xa, or 250nM thrombin. Quantification of the ratio of green fluorescent area to total confluence. (4 fields/replicate well, 4 wells/condition). (**B**) Nanoluciferase luminescent signal was measured following infection with rVSVΔG/SARS-CoV-2 pseudovirus and the addition of either vehicle, factor Xa, or thrombin. The effect of factor Xa on rVSVΔG complemented with either (**C**) SARS-CoV spike or (**D**) VSV-G was measured by luminescent signal. Luminescent signal was measured following HIV-1_NL_/SARS-CoV-2 pseudovirus infection and concomitant treatment with 125-250nM factor Xa in (**E**) Calu3 cells and (**F**) A549/ACE2 and. (**G**) Vero cells were transduced with lentiviral vectors to express GFP or TMPRSS2. Following selection, cells were infected with HIV-1_NL_/SARS-CoV-2 pseudovirus and concomitantly treated with 125-250nM factor Xa. Subsequently, Nanoluciferase luminescent signal was determined and plotted relative to vehicle-treated control. * P<0.05, two-tailed t-test. Error bars represent +/− SEM.

While factor X and prothrombin levels are variable between individuals in healthy populations (Brummel-Ziedins, Orfeo, Gissel, Mann, & Rosendaal, 2012), the concentration of proteases used in the pseudovirus assay (125-250nM) are comparable to reference ranges of factor X (Brummel-Ziedins et al., 2012; Tormoen, Khader, Gruber, & McCarty, 2013; Williams & Marks, 1994) and prothrombin (Baugh, Broze, & Krishnaswamy, 1998; Brummel-Ziedins et al., 2012; Tormoen et al., 2013). Similar concentrations of active purified proteases were required to normalize *in vitro* clotting times, where purified factor Xa was used to correct dilute Russell’s viper venom time (dRVVT) of factor X-deficient human plasma **(Fig. S7A)** and purified thrombin was used to correct the prothrombin time of prothrombin-deficient human plasma **(Fig. S7B)**.

SARS-CoV-2 contains a notable insertion of basic residues at the S1/S2 boundary, distinguishing its sequence from many related betacoronaviruses (Jaimes, Andre, Chappie, Millet, & Whittaker, 2020). Entry of rVSVΔG was increased when complemented with spike protein from SARS-CoV of the 2002 outbreak (**Fig. 3C**), but not when complemented instead with VSV G (**Fig. 3D**). This indicates that factor Xa spike cleavage could be relevant across multiple coronaviruses, but is not generally associated with VSV entry.

We further validated that factor Xa activated spike-mediated entry the HIV-1-based pseudovirus system (Schmidt et al., 2020). Consistent with the results above, addition of purified Factor Xa to the media at the time of infection enhanced entry of HIV-1-based SARS-CoV-2 pseudovirus in Calu3 cells (**Fig. 3E**). Thrombin did not appear to enhance spike-mediated entry by HIV-1/SARS-CoV-2 pseudovirus, unlike rVSVΔG/SARS-CoV-2 pseudovirus (**Fig. S6B-E**).

We investigated the functional interplay of TMPRSS2 expression and the effect of exogenous activated coagulation factors. TMPRSS2 is expressed in Calu3 cells and contributes to coronavirus entry (Hoffmann, Kleine-Weber, et al., 2020), whereas A549/ACE2 and Vero cells lack endogenous TMPRSS2 expression. Factor Xa induced a significant dose-dependent effect on pseudovirus entry in both Calu3 and A549/ACE2 (**Fig. 3E-F**). Furthermore, an isogenic pair of Vero cells was generated by expressing TMPRSS2 or GFP control. Pseudovirus infection of both Vero^GFP^ and Vero^TMPRSS2^ cells were significantly increased by factor Xa, indicating that factor Xa enhancement of infection is not dependent on TMPRSS2 (**Fig. 3G**). This is consistent with the model that FXa cuts the S1/S2 site and TMPRSS2 has functionally important activity at the S2’ site.

### Nafamostat broadly inhibits cleavage of spike peptides by both transmembrane serine proteases and coagulation factors

Given that multiple proteases could contribute to SARS-CoV-2 spike cleavage activation, a drug that could inhibit both transmembrane serine proteases and coagulation factors would be potentially valuable as a dual action anticoagulant/antiviral in COVID-19. Accordingly, we next explored the candidate set of inhibitors for cross-reactivity against a broader set of proteases that could facilitate viral entry. Like TMPRSS2, human airway trypsin-like protease (HAT), encoded by *TMPRSS11D*, is a member of the TTSP family of proteases and could activate SARS-CoV-2 S (Hoffmann et al., 2021). HAT exhibited sensitivity to camostat and nafamostat, similar to TMPRSS2 (**Fig. 4A-B**). Compared to TMPRSS2, HAT was more sensitive to dabigatran and less sensitive to otamixaban (**Fig. 4A-B**). Factor Xa activity against the S1/S2 peptide was most sensitive to otamixaban and moderately sensitive to nafamostat and dabigatran (**Fig. 4C)**. Thrombin activity was sensitive to camostat, nafamostat, and dabigatran, and moderately sensitive to otamixaban (**Fig. 4D**).

**Figure 4.**
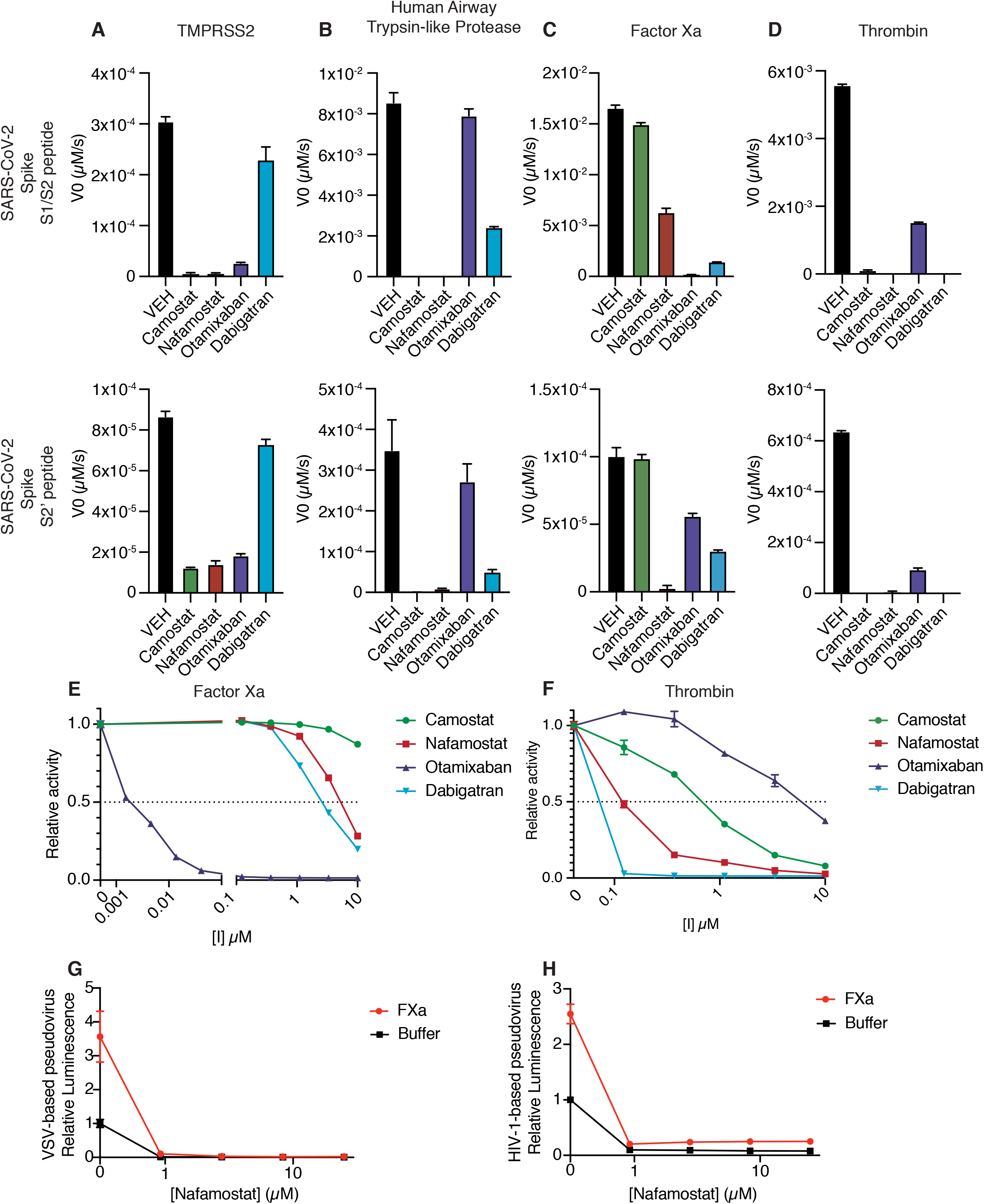
Nafamostat broadly inhibits cleavage of spike peptides by both coagulation factors and transmembrane serine proteases. Initial velocities for the cleavage of 10 *µ*M SARS-CoV-2 spike S1/S2 (top) and S2’ (bottom) peptide substrates by (**A**) TMPRSS2, (**B**) TMPRSS11D/Human airway trypsin-like protease (**C**) factor Xa, and (**D**) thrombin were measured in the presence of DMSO vehicle, or 10*µ*M camostat, nafamostat, otamixaban, or dabigatran. The relative activity of (**E**) factor Xa and (**F**) thrombin were determined over a range of 0-10*µ*M of the indicated drugs. Calu3 cells were treated with a range of concentrations of nafamostat with or without addition of 250 nM exogenous factor Xa and infected with (**G**) rVSVΔG/SARS-CoV-2 pseudovirus or (**H**) HIV-1_NL_/SARS-CoV-2 pseudovirus and infectivify was measured by luminescence. Error bars +/− SEM.

We performed a dose response curve of the panel of inhibitors on factor Xa and thrombin S1/S2 cleavage. Otamixaban, a designed factor Xa inhibitor, demonstrated an IC50 at the nanomolar level to factor Xa, while nafamostat and dabigatran demonstrated IC50’s in the micromolar range (**Fig. 4E**). Camostat did not potently inhibit factor Xa spike cleavage. Dabigatran, a designed thrombin inhibitor, as well as nafamostat and camostat demonstrated a submicromolar IC50 for thrombin-dependent spike cleavage (**Fig. 4F**). Otamixaban inhibited thrombin spike cleavage in the micromolar range.

Furin showed high activity against the S1/S2 peptide, but not against the S2’ peptide, and was not sensitive to any of the candidate inhibitors (**Fig. S5A-B**). While it has been suggested that TMPRSS4 or neutrophil elastase may also cleave SARS-CoV-2 spike, we detected minimal activity against either S1/S2 or S2’ peptide substrates in our enzymatic assay (<1% of furin cleavage of S1/S2) (**Fig. S5A-B**).

In the pseudovirus assay, nafamostat effectively suppresses SARS-CoV-2 S-mediated entry with or without the addition of exogenous factor Xa, using either the VSV-based (**Fig. 4G**) or HIV-1-based (**Fig. 4H**) SARS-CoV-2 pseudovirus. To clarify the pleiotropic nature of nafamostat, which inhibits TMPRSS2 and factor Xa, we compared the effect of apixaban, which inhibits factor Xa but not TMPRSS2. Apixaban rescued the effect of exogenous FXa back to the baseline level of infection, but did not affect pseudovirus infection in the absence of exogenous protease (**Fig. S9**). Direct oral anticoagulants (DOACs) have the potential to block clotting factor-mediated enhancement of viral entry, but TMPRSS2-mediated cleavage activation would remain unaffected by treatment with DOACs in common usage in North America and Europe. Taken together, nafamostat appears to be a versatile inhibitor of spike activation by a variety of TTSPs and coagulation factors. The multitarget mechanism of nafamostat distinguishes its potential as an antiviral/anticoagulant from currently FDA-approved DOACs.

## Discussion

### Coagulation factors cleave the SARS-CoV-2 spike protein

Using a FRET-based enzymatic assay and two platforms of pseudovirus assays, we demonstrate that coagulation proteases factor Xa and thrombin cleave SARS-CoV-2 spike protein. Coagulation-induced cleavage enhances spike activation and increases viral entry into target cells, potentially instigating a positive feedback loop with infection-induced coagulation. Nafamostat, among currently available drugs, is best suited as a multi-purpose inhibitor against spike cleavage by TTSPs and coagulation factors. These data have numerous implications at the intersection of virology and coagulation.

### Viral envelope protein activation by non-target-cell proteases

Hijacking of host transmembrane, endosomal, and ER proteases to activate viral envelope proteins has been described in influenza A, human metapneumovirus, HIV, and Sendai virus (Kido, Niwa, Beppu, & Towatari, 1996; Straus, Kinder, Segall, Dutch, & Whittaker, 2020). In the present study, we find an instance where the virus can be primed not by proteases expressed by the target cell, but by host organism proteases derived from the microenvironment of the target cell. Prior studies have described cleavage activation of SARS-CoV by elastase and plasmin, illustrating that microenvironmental host proteases can indeed play an important role in coronavirus spike priming (Belouzard, Madu, & Whittaker, 2010; Kam et al., 2009; Matsuyama, Ujike, Morikawa, Tashiro, & Taguchi, 2005). Generally, the scope by which circulating proteases, such as coagulation factors, or proteases expressed by immune cells interact with viral envelope proteins during infection has not been comprehensively explored.

Relatively few studies have examined the interaction of factor Xa or thrombin and viral proteins, and each relies on target cells as the source of coagulation factors. Our results are consistent with a prior study that concluded that factor Xa cleaves and activates SARS-CoV spike (Du et al., 2007). Traditionally, influenza vaccines rely on viral propagation in chicken eggs, where hemagglutinin cleavability by factor Xa is a determinant of the efficiency of strain-specific propagation of influenza A virus *in ovo* (Gotoh et al., 1990; Straus & Whittaker, 2017). Hepatitis E virus ORF1 polyprotein is processed intracellularly by thrombin and factor Xa in the cytoplasm of hepatocytes, which are the primary cell type responsible for generating and secreting coagulation factors (Kanade, Pingale, & Karpe, 2018).

Activation of coagulation has the potential to exacerbate SARS-CoV-2 infectivity in both TMPRSS2+ and TMPRSS2-host cells. Reliance on extracellular proteolytic activity could expand the field of susceptible cell types and regions of the airway. Extrapulmonary infection has been described, particularly in small intestinal enterocytes (Xiao et al., 2020; Zang et al., 2020) and, in some cases, the central nervous system (Song et al., 2021). It warrants investigation whether hypercoagulation is linked to extrapulmonary infection.

### Evolutionary perspective on viral-host interaction

Proteolytic cleavage of the spike forms a barrier to zoonotic crossover independent of receptor binding (Menachery et al., 2020). Hemostasis is of central importance in mammals and represents a major vulnerability of mammals to predators and pathogens, either through hyperactivation of coagulation or uncontrolled bleeding. The dysregulation of hemostasis is a convergent mechanism of toxins of snakes, bees, and bats (Ma et al., 2013; Markland & Swenson, 2013; Prado, Solano-Trejos, & Lomonte, 2010) and a driver of virulence in Ebola and dengue virus infection (Geisbert et al., 2003; Rathore et al., 2019). Acute lung injury from viral cytopathic effects, the induction of a COVID-19-associated cytokine storm, complement activation, and anti-phospholipid autoantibodies have all been suggested to instigate the coagulation cascade (Merrill, Erkan, Winakur, & James, 2020; Zuo et al., 2020). Perhaps, SARS-CoV-2 has undergone selection to exploit an environment locally enriched in coagulation proteases for enhanced entry. As infection spreads, more clotting is induced, instigating a positive feedback loop to promote entry into additional host cells.

### Clinical relevance of potential antiviral activity of anticoagulants

Effective anticoagulation is a critical area of investigation to improve outcomes in coronavirus infection. Vitamin K antagonists, including heparin, are commonly used for preventing venous thromboembolism in COVID-19, although no strong evidence yet supports any specific anticoagulant (Cuker et al., 2021). Three large randomized clinical trials to determine the benefit of therapeutic-intensity vs. prophylactic intensity heparin in critically ill COVID-19 patients were suspended at interim analysis for futility (NCT02735707, NCT04505774, and NCT04372589). There has been interest in the use of direct-acting oral anticoagulants (DOACs) to manage COVID-19 related coagulopathy, but optimal protocols for managing coagulopathy in COVID-19 patients have not yet been developed (Capell et al., 2021; Lopes et al., 2021). The most prominent risk of anticoagulants is bleeding, and notably direct oral anticoagulants, as well as nafamostat, have a reduced risk of intracranial hemorrhage and other bleeding events compared to vitamin K antagonists (Chen, Stecker, & B, 2020; Hellenbart, Faulkenberg, & Finks, 2017; Makino et al., 2016).

In our studies, anticoagulant serine protease inhibitors, otamixaban and dabigatran, exhibited off-target activity against TMPRSS2 and other TTSPs, but likely require concentrations higher than those safely reached *in vivo* (Paccaly et al., 2006; Stangier & Clemens, 2009). On the other hand, our data suggest that nafamostat and camostat may offer three distinct therapeutic mechanisms against SARS-CoV-2 infection; these compounds have the potential to block spike cleavage mediated by TMPRSS2 and other TTSPs, to block spike cleavage by coagulation factors, and to serve as an anticoagulant. Nafamostat (Fujii & Hitomi, 1981; Keck et al., 2001; Takeda, Matsuno, Sunamura, & Kakugawa, 1996) and Camostat (Ramsey, Nuttall, Hart, & Team, 2019) have been in clinical use in Asia for many years for the treatment of pancreatitis. Nafamostat has also been used as an anticoagulant during hemodialysis (Akizawa et al., 1993) and extracorporeal membrane oxygenation (ECMO) (Park et al., 2015), and to manage disseminated intravascular coagulopathy (Kobayashi, Terao, Maki, & Ikenoue, 2001). As of this writing, there are currently eight open clinical trials listed on clinicaltrials.gov to investigate the use of nafamostat in COVID-19, while 23 active clinical trials of camostat for COVID-19 were identified.

Inhibition of coagulation factor-induced spike cleavage may contribute to the molecular mechanism of these agents, if treatment is given sufficiently early. Many COVID-19 associated complications leading to hospitalization occur as immune hyperactivation waxes and peak viral titer wanes (Griffin et al., 2021). To take advantage of the potential antiviral effect of anticoagulants, early intervention in an outpatient setting may be beneficial.

### Limitations

The experiments of this study, like prior studies using similar techniques, have some limitations. Protease enzymatic assays on peptide substrates allow for detailed biochemical characterization of a specific site, but peptide substrates may not have the equivalent three-dimensional conformation or post-translational modifications of the full-length protein produced in appropriate cells. For instance, SARS-CoV-2 S is extensively glycosylated (Watanabe, Allen, Wrapp, McLellan, & Crispin, 2020). The possibility of additional spike cleavage sites and potential pro- and anti-viral consequence of proteases acting on cell surface proteins including ACE2 cannot be excluded. The amount, density, and accessibility of spike protein could be different between these cell-based surrogate assays and wild type SARS-CoV-2 infection. However, antibody neutralization is highly correlated between authentic virus and corresponding pseudotyped viruses, suggesting similar conformation (Schmidt et al., 2020). We have attempted to mitigated the risk of artifact by using multiple orthogonal platforms.

### Conclusion

Collectively, our data provide rationale for the investigation of early intervention with judiciously selected anticoagulant treatment, which may have collateral benefit in limiting progressive spread of SARS-CoV-2 infection throughout the lung in infected individuals. Preparedness to mitigate a future SARS-CoV-3 epidemic is critical to pursue through the understanding of coronavirus-host interactions.

## Author Contributions

Conceptualization, E.R.K and L.C.C.; Methodology, E.R.K, J.A.J., J.L.J., F.M., Y.W.; Investigation, E.R.K., J.A.J., M.M, A.E.S., Y.B.; Writing – Original Draft, E.R.K. and L.C.C.; Writing – Review & Editing, E.R.K., J.L.J., F.M., Y.B., R.E.S., G.W., and L.C.C.; Funding Acquisition, R.E.S., G.W., and L.C.C.; Resources, F.M., Y.W.; Supervision, R.E.S., G.W., and L.C.C..

## Acknowledgements

The authors would like to thank Pilar Mendoza (Rockefeller University), John Blenis (WCMC), Elena Piskounova (WCMC), Tim McGraw (WCMC), Marco Straus (Cornell University), Tomer Yaron (WCMC), and all members of the Cantley Lab for insightful discussion and helpful comments. We thank Paul Bieniasz (Rockefeller University) and Theodora Hatziioannou (Rockefeller University) for providing reagents and helping to establish the pseudovirus platform in our laboratory. We thank Danielle Bulaon (Weill Cornell) for provisioning inhibitors, Benjamin tenOever (Mount Sinai) for providing Vero cells, and Francisco Sanchez-Rivera and Scott Lowe (MSKCC) for the ipUSEPR plasmid. This work was funded in part by the National Institute of Health research grant R01AI35270 (to GW).

## Supplementary Data

**Supplementary Figure 1.**
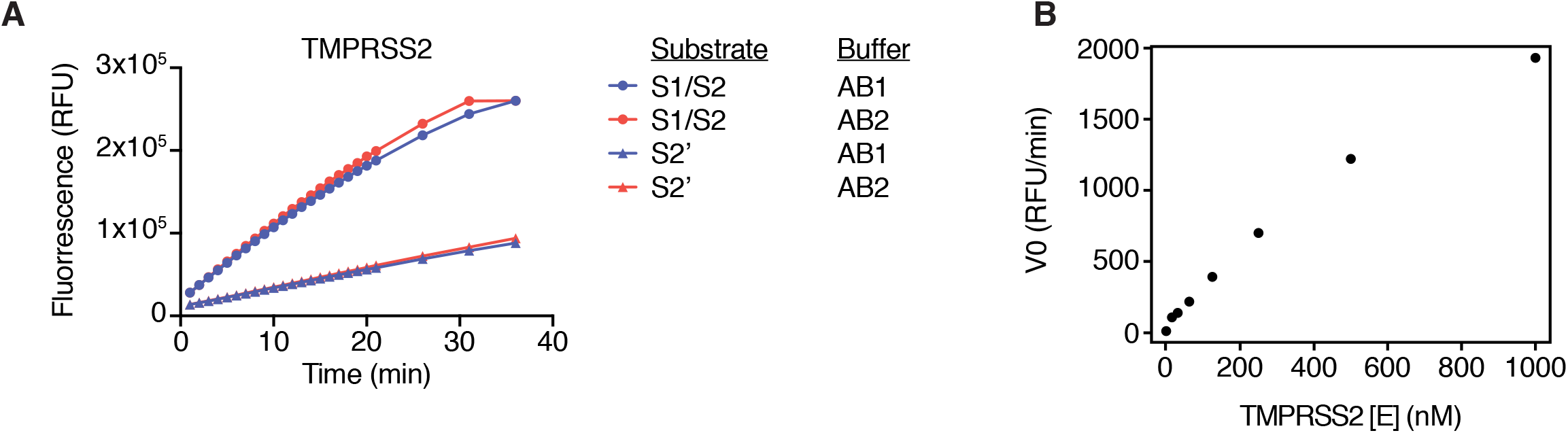
Optimization of FRET enzymatic assay, related to Figure 1. (**A**) TMPRSS2 enzymatic assay was performed in AB1 (20mM Tris-HCl, pH7.3, 100 mM NaCl, 1 mM EDTA, fresh 1 mM DTT) or AB2 (50mM Tris-HCl, 150mM NaCl, pH 8) using 10 *µ*M of either S1/S2 or S2’ peptide substrate. (**B**) Titration of enzyme concentration was performed (0-1000 nM) with 10 *µ*M S1/S2 substrate. Initial reaction velocity V_0_ (rate of change in fluorescent signal) each enzyme concentration with 10*µ*M S1/S2 peptide substrate.

**Supplementary Figure 2.**
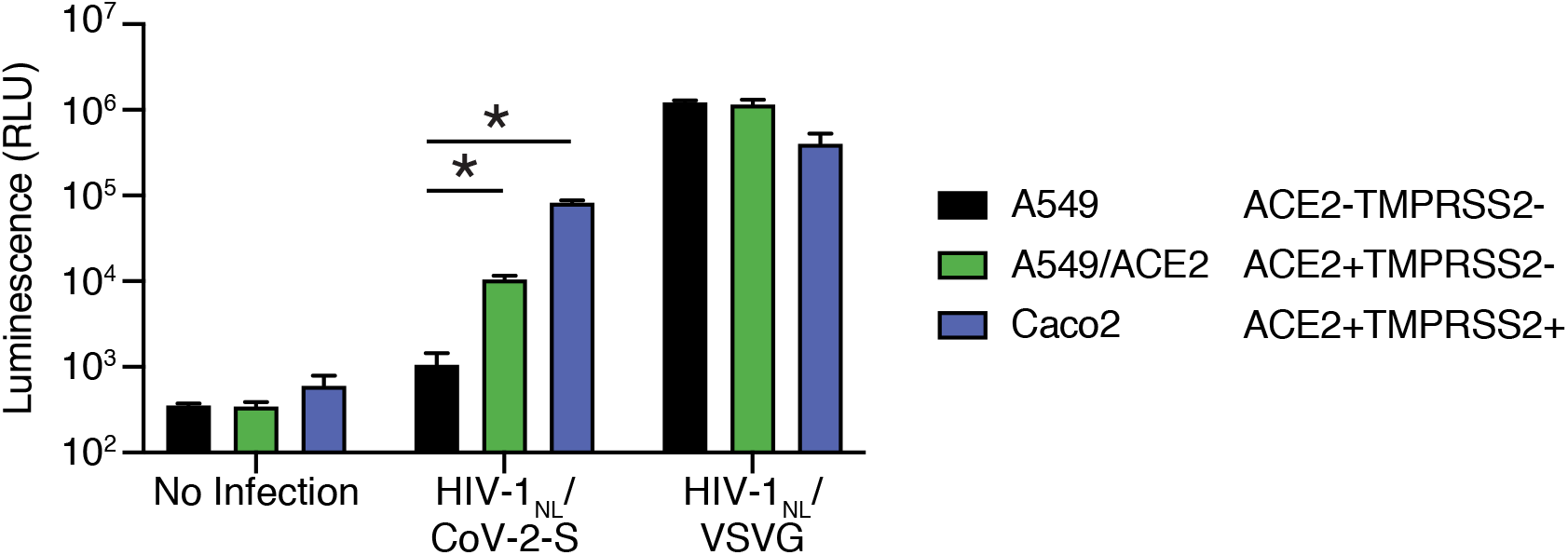
Further characterization of HIV-1/SARS-CoV-2 pseudovirus, related to Figure 1. A549 cells (which do not express ACE2), A549/ACE2 cells (ectopic ACE2 expression from a lentiviral vector), and Caco2 cells (which express endogenous ACE2 and TMPRSS2) infected with HIV-1_NL_-based particles pseudotyped with SARS-CoV-2 S or VSV G. * P<0.05, two-tailed t-test. Error bars +/− SEM.

**Supplementary Figure 3.**
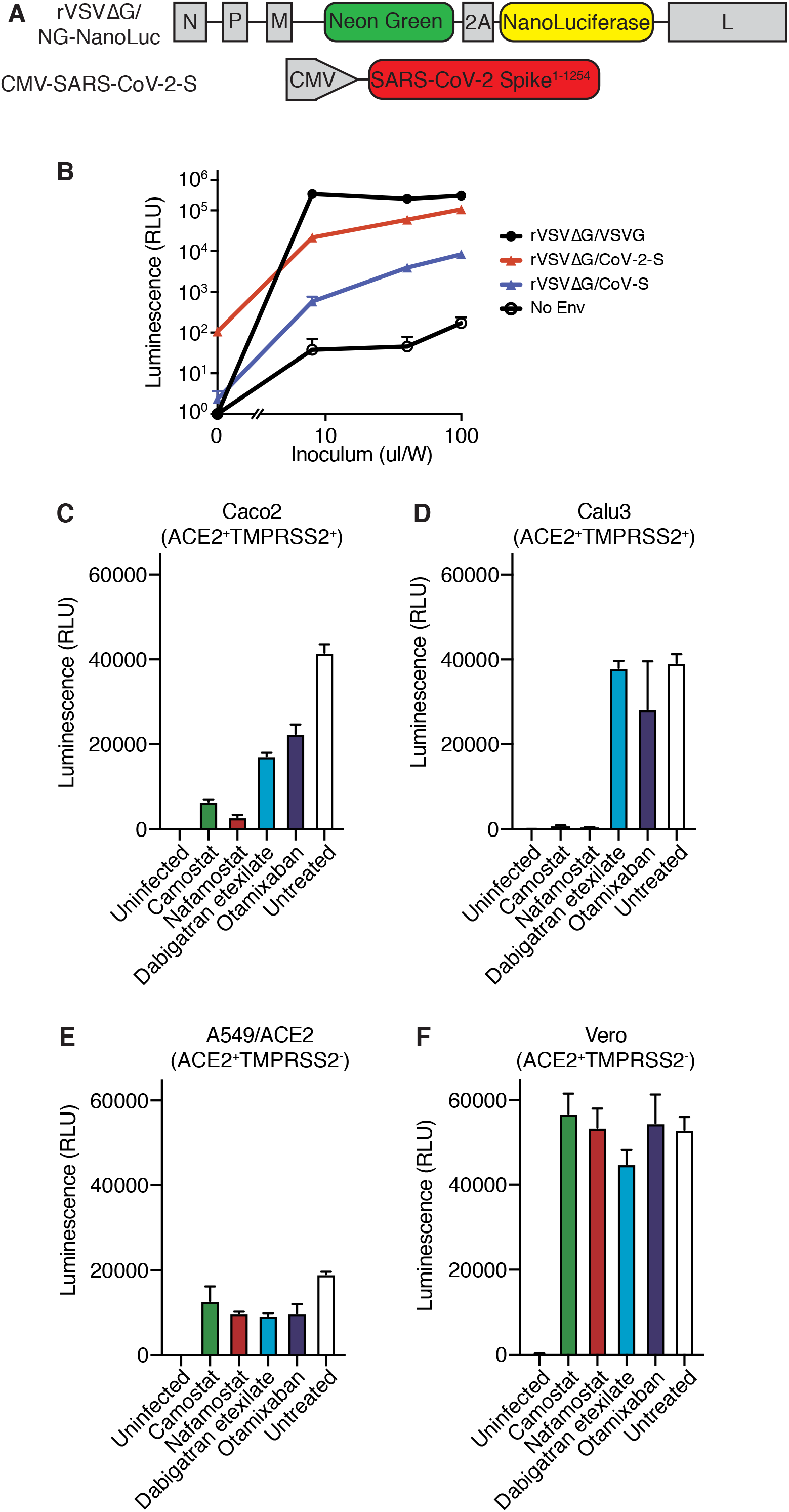
Further characterization of rVSVΔG/SARS-CoV-2 pseudovirus, related to Figure 1. (**A**) Schematic of constructs used to generate SARS-CoV-2 spike-pseudotyped/VSV-based pseudovirus. (**B**) Nanoluciferase luminescent signal following addition of rVSVΔG pseudovirus complemented with VSV G, SARS-CoV-2 S, SARS-CoV S, or without complementation with any envelope protein to Calu3 cells. Each pseudovirus was titrated by adding the indicated volume of inoculum, supplemented with fresh media up to 200 ul/well in a 96W plate. (**C-F**) Nanoluciferase luminescent signal following infection of (**C**) Caco2 (**D**) Calu3, (**E**) A549/ACE2, or (**F**) Vero cells with rVSVΔG/SARS-CoV-2 pseudovirus pretreated for 4 hours with 10 *µ*M camostat, nafamostat, dabigatran, or otamixaban, compared with uninfected or infected/untreated cells. Expression status of ACE2 and TMPRSS2 for each cell line is indicated.

**Supplementary Figure 4.**
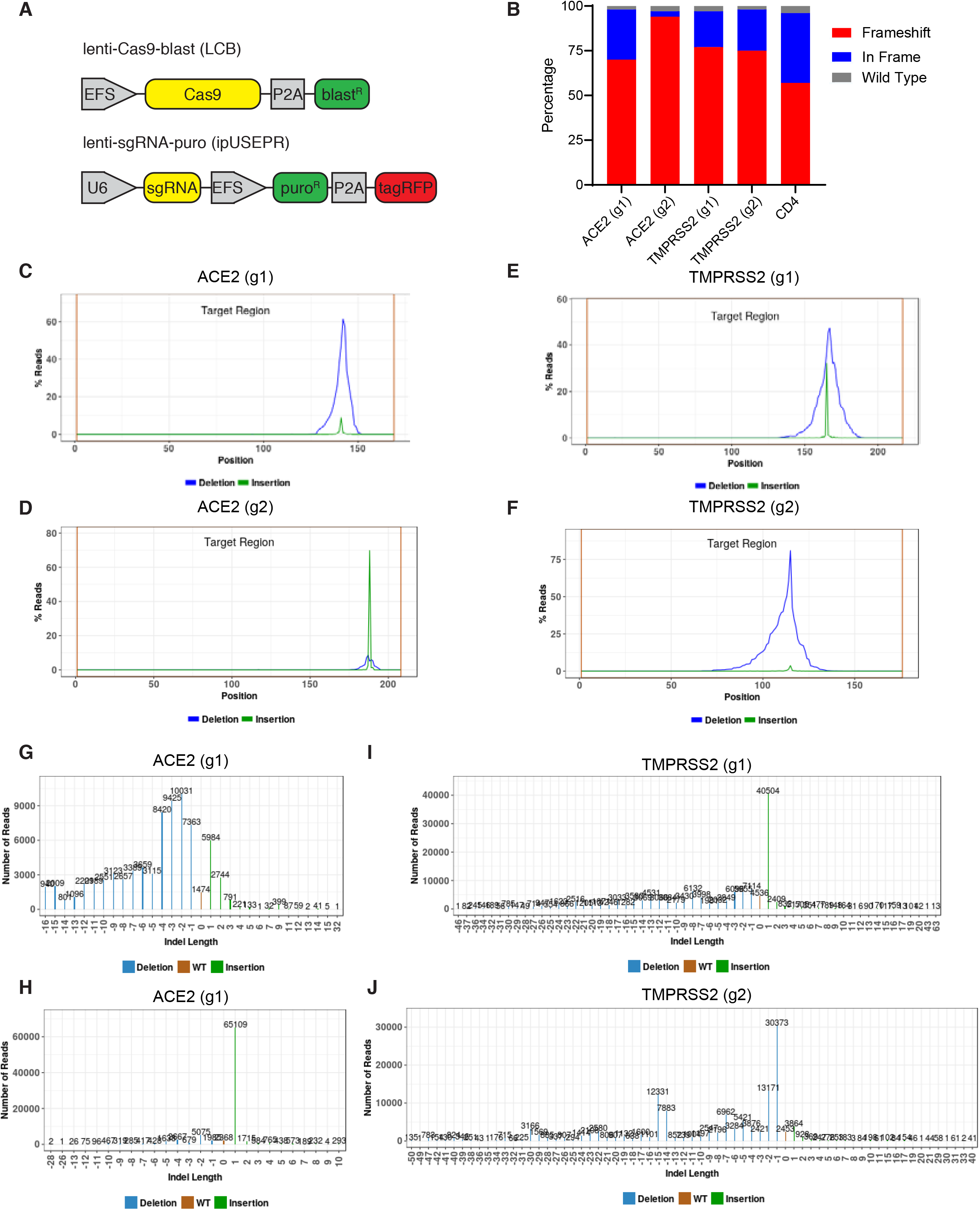
Evidence of CRISPR knockout efficiency, related to Figure 1. (**A**) Constructs used for CRISPR experiments. (**B**) Percentage of reads exhibiting wild type, frameshift, or in-frame indels at each locus for the indicated sgRNAs. (**C-F**) Distribution of reads with deletion or insertion by position within amplicon. (**G-J**) Distribution of the size of insertions and deletions in each amplicon. Two sgRNAs targeting ACE2 (g1 and g2) and two sgRNAs targeting TMPRSS2 (g1 and g2) were analyzed.

**Supplementary Figure 5.**
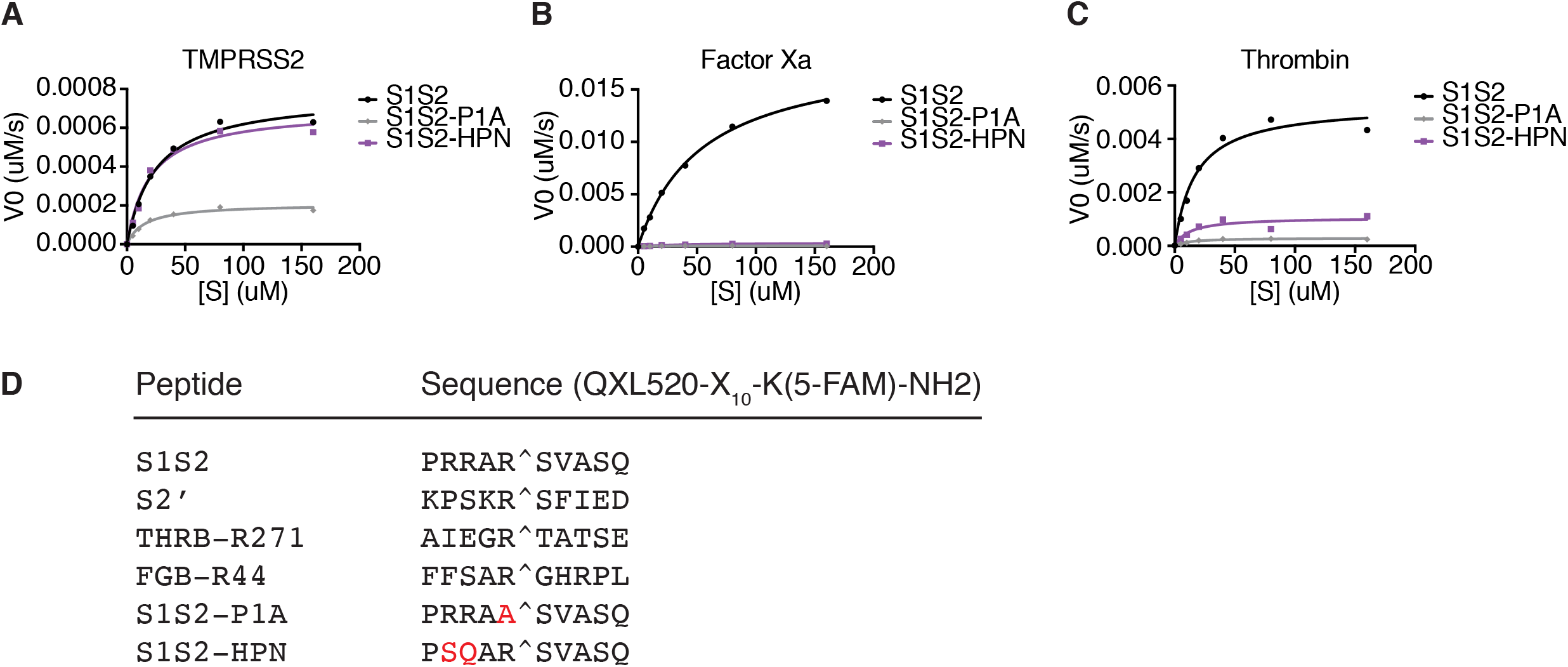
FRET enzymatic assay with modified peptide substrates, related to Figure 2. (**A-C**) Initial reaction velocity with respect to enzyme concentration for peptide substrates of the SARS-CoV-2 spike S1/S2 site (S1S2), with P1 arginine substituted with alanine (S1S2-P1A), or with substitutions in the P3 and P4 position (RR>SQ) with (**A**) TMPRSS2, (**B**) factor Xa, or (**C**) thrombin. (**D**) List of peptide substrates used in this study.

**Supplementary Figure 6.**
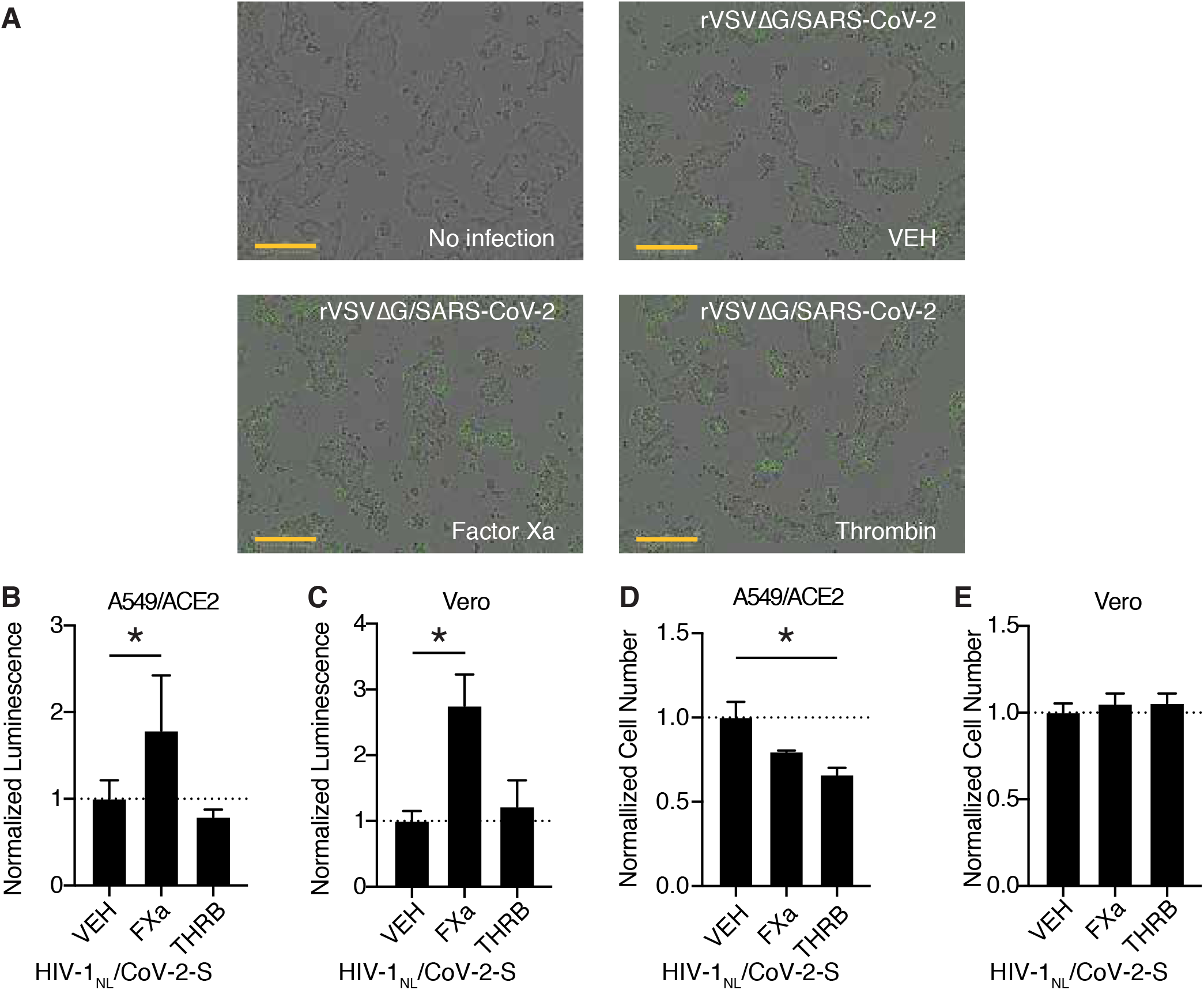
Further characterization of coagulation factor-induced SARS-CoV-2 pseudovirus infection, related to Figure 3. (**A**) Representative merged brightfield and green fluorescence images of Calu3 cells without infection or following rVSVΔG/SARS-CoV-2 pseudovirus infection and concomitant treatment with vehicle, 250nM factor Xa, or 250nM thrombin (corresponding to **Fig. 3A**). Scale bars represent 300 *µ*m. (**B-E**) HIV-1_NL_/SARS-CoV-2 pseudovirus with addition of purified protease or vehicle. Nanoluciferase luminescent signal relative to vehicle-treated control was measured following infection in (**B**) A549/ACE2 and (**C**) Vero cells. Cell number following protease treatment, relative to vehicle control was determined for (**D**) A549/ACE2 and (**E**) Vero cells. * P<0.05, two-tailed t-test. Error bars +/− SEM.

**Supplementary Figure 7.**
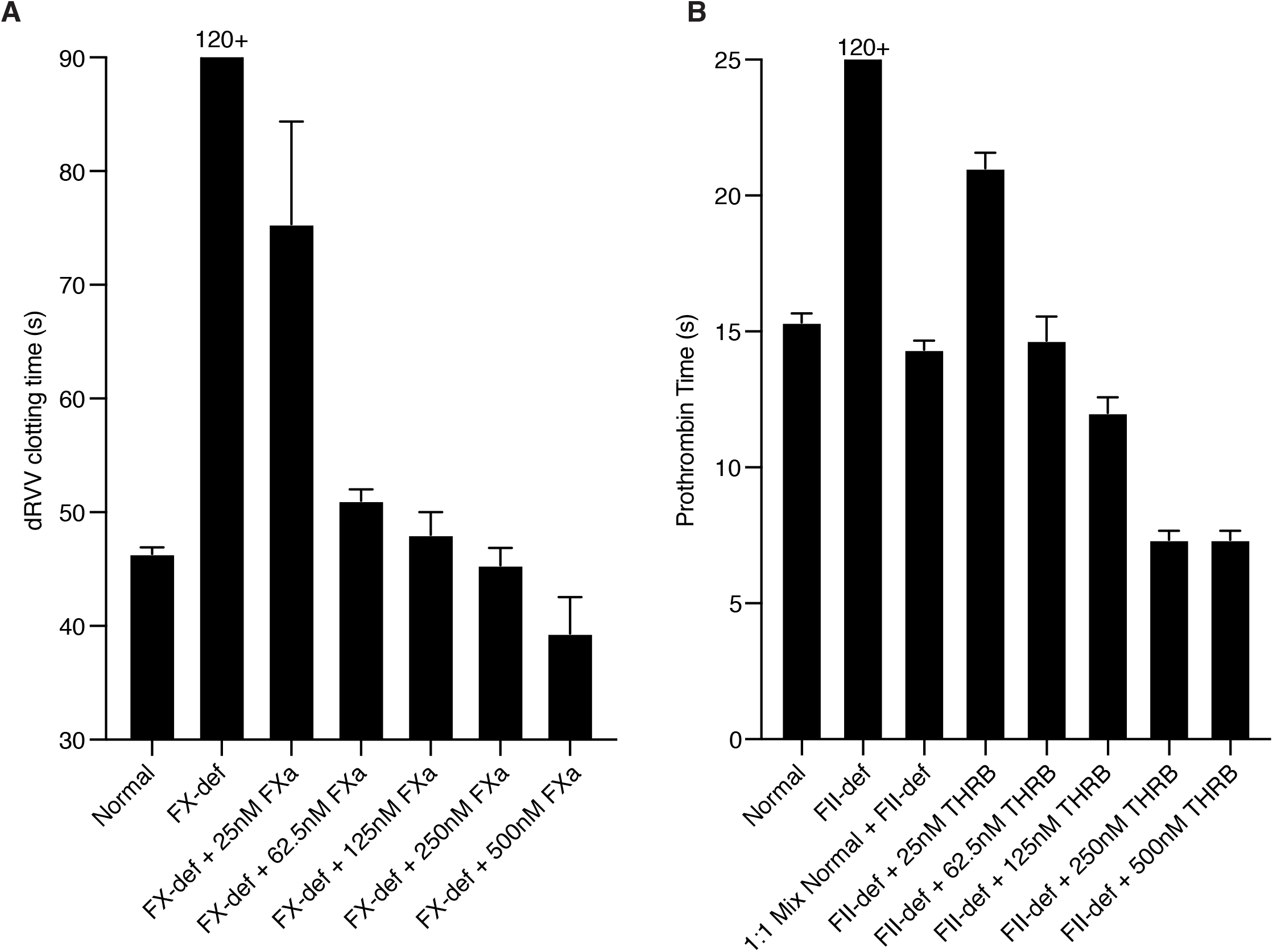
Assessing relevant protease levels with *ex vivo* clotting assays, related to Figure 3. (**A**) Factor X activity was determined by dilute Russell’s Viper Venom clotting time (dRVVT). Pooled normal plasma or FX-deficient plasma with the addition of the indicated concentration of active purified factor Xa were assayed. (**B**) Prothrombin activity was determined by prothrombin time (PT) via activation with thromboplastin. Pooled normal plasma or prothrombin-deficient plasma with the addition of the indicated concentration of active purified thrombin were assayed.

**Supplementary Figure 8.**
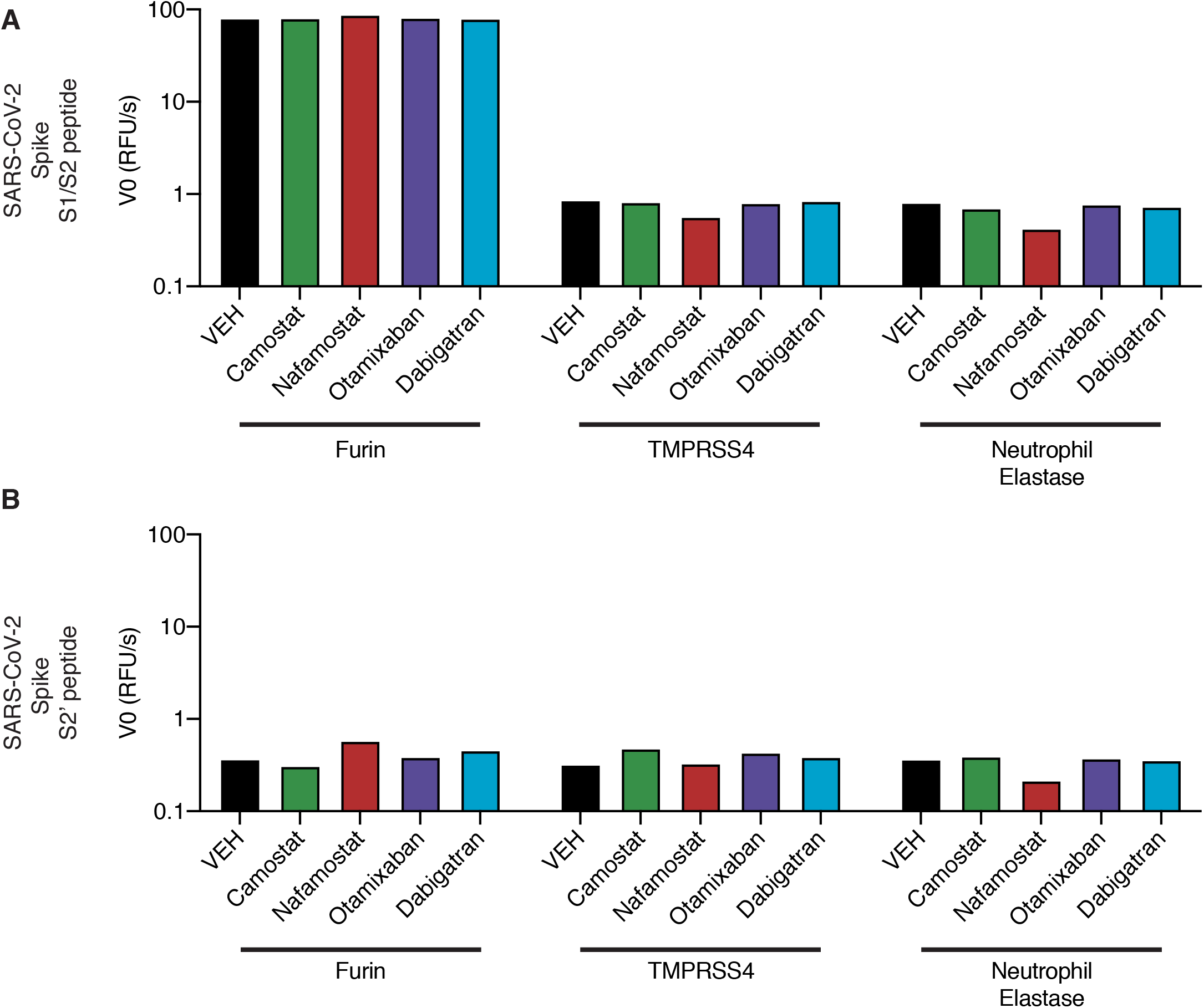
Activity of candidate inhibitors against other proteases, related to Figure 4. Initial reaction velocity V_0_ of furin, TMPRSS4, or neutrophil elastase cleavage of (**A**) S1/S2 peptide substrate or (**B**) S2’ peptide substrate, treated with DMSO vehicle, camostat, nafamostat, otamixaban, or dabigatran. 10*µ*M Substrate and 10*µ*M inhibitor were used.

**Supplementary Figure 9.**
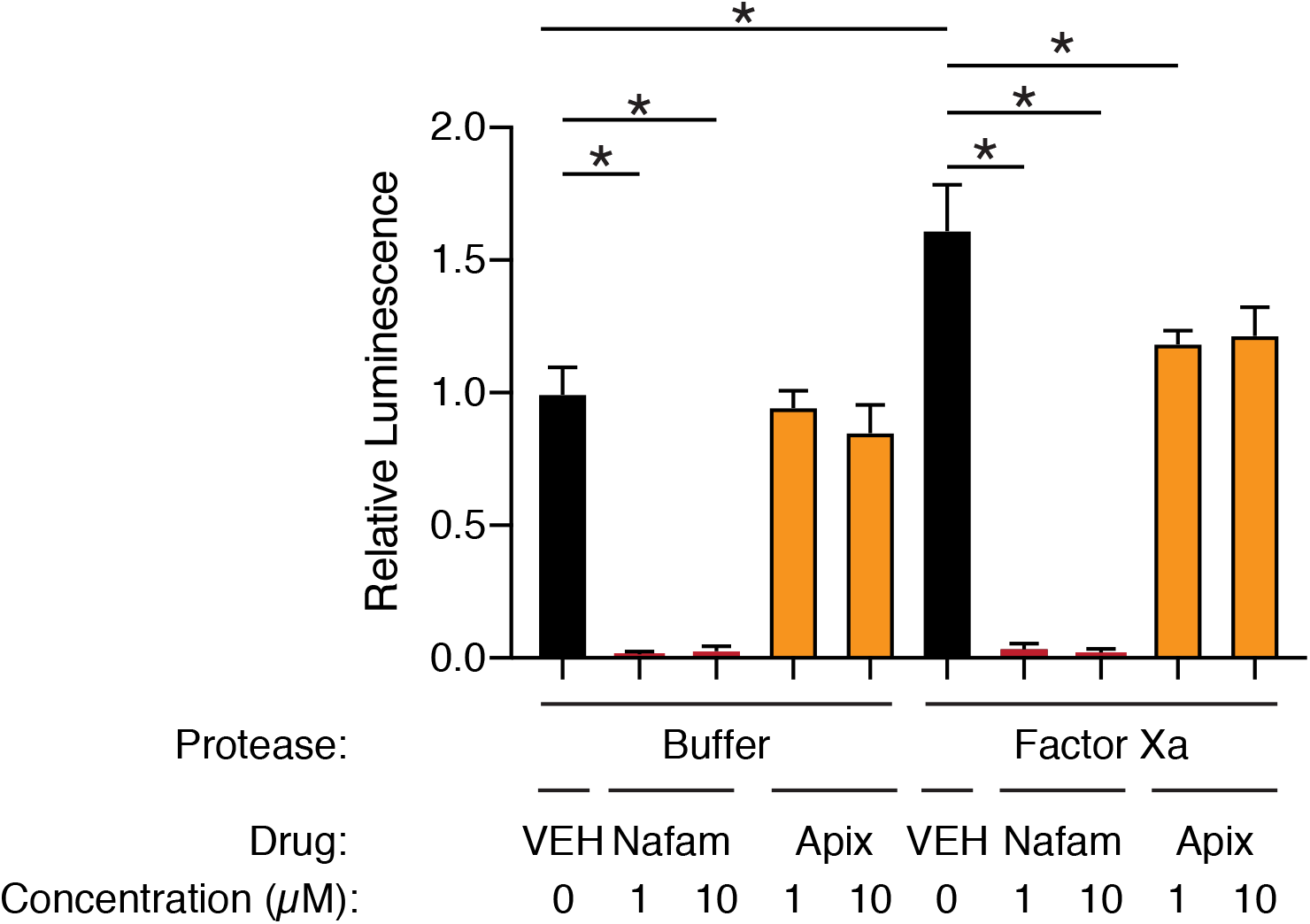
Apixaban rescues effect of factor Xa, related to Figure 4. Calu3 cells were infected with rVSVΔG/SARS-CoV-2 pseudovirus with addition of protease buffer (left) or factor Xa (right). Cells were treated at the time of infection with DMSO vehicle (black), 1 *µ*M or 10 *µ*M nafamostat (red), or 1 *µ*M or 10 *µ*M apixaban (orange). * P<0.05, two-tailed t-test. Error bars +/− SEM.

## Methods

### Enzymatic Assay

Thrombin (605195, human, activated by factor Xa, factor Va, and phospholipid) and factor Xa (69036, bovine, activated by Russell’s Viper Venom), were obtained from Millipore Sigma. Recombinant TMPRSS2, purified from yeast, was obtained from LSBio (LS-G57269). TMPRSS4 was obtained from Aviva System Biology (OPCA0240), furin was obtained from Thermo Fisher Scientific (1503SE010), neutrophil elastase was obtained from Thermo Fisher Scientific (9167SE020). FRET peptides were obtained from Anaspec and a peptide sequences are listed in **Supplementary Fig S1F**. Protease assay buffer was composed of 50mM Tris-HCl, 150mM NaCl, pH 8. Enzyme dilution/storage buffer was 20mM Tris-HCl, 500mM NaCl, 2mM CaCl_2_, 50% glycerol, pH 8. Peptides were reconstituted and diluted in DMSO. Enzyme kinetics were assayed in black 96W plates with clear bottom and measured using a BMG Labtech FLUOstar Omega plate reader, reading fluorescence (excitation 485nm, emission 520nm) every minute for 20 cycles, followed by every 5 minutes for an additional 8 cycles. A standard curve of 5-FAM from 0-10 *µ*M (1:2 serial dilutions) was used to convert RFU to *µ*M of cleaved FRET peptide product. Calculation of enzyme constants was performed with Graphpad Prism software (version 9.0). Camostat and nafamostat were obtained from Selleck Chemicals and all other inhibitors were obtained from MedChem Express.

### Cell Culture

Calu3, A549, Caco2, and Vero cells were tested for mycoplasma (Lonza MycoAlert detection kit) and human cell line identity was authenticated by ATCC. A549 and Vero cells were grown in DMEM, supplemented with 10% FBS, 100U/ml Penicillin, and 100ug/ml Streptomycin. Calu3 and Caco2 cells were grown in MEM, supplemented with 10% FBS, 100U/ml Penicillin, 100ug/ml Streptomycin, 1% MEM NEAA and 1mM sodium pyruvate.

### Plasmids and lentivirus infection

Overexpression constructs pEGPN-GFP, pEGPN-ACE2, and pEGPN-TMPRSS2 were constructed by Gibson cloning using NEBuilder master mix (New England Biolabs, E2621) with overlapping PCR generated inserts for promoter EF1*α*, the gene of interest, promoter PGK, and neomycin/resistance gene. Lentiviral vectors were co-transduced with MD2G and PAX2 in 293T cells (5 million cells/10cm plate) with 25ul of XtremeGene9 (Millipore Sigma, #6365787001) and supernatant was harvested at 48hr and 72hr post transfection. Target cells were transduced with the addition of 4 *µ*g/ml polybrene (Santa Cruz, sc-134220). Infected cells were selected and maintained in 500 ug/ml G418 (Life Technologies, #10131027). lentiCas9-Blast was a gift from Feng Zhang (Sanjana, Shalem, & Zhang, 2014) (Addgene plasmid # 52962). ipUSEPR was a gift from Francisco Sanchez-Rivera and Scott Lowe. sgRNAs were selected from the Brunello CRISPR database (Doench et al., 2016). Four guides per gene were tested in Caco2 cells and the most efficient two sgRNAs/gene were used in subsequent experiments (**Supplementary Fig. S4**). Knockout efficiency was determined by next-generation amplicon sequencing (Genewiz).

### Pseudovirus

Recombinant VSV-based and HIV-1-based SARS-CoV-2 pseudovirus was generated as described previously (Schmidt et al., 2020). To generate rVSVΔG/SARS-CoV-2 pseudovirus, 293T cells (12 million cells/15cm plate) were transfected with 12.5 *µ*g pSARS-CoV-2_Δ19_, and 24 hr post-transfection, were infected with VSV-G-complemented rVSVΔG virus at an MOI of 1. Supernatant was collected 16 hr post-infection, centrifuged at 350 g x 10min, filtered through a 0.45-*µ*m filter, and concentrated using Lenti-X-Concentrator (Takara Bio). Prior to infection of target cells, the viral stock was incubated with anti-VSV-G antibody (3ug/ml) for 1 h at 37°C to neutralize contaminating rVSVΔG/NG/NanoLuc/VSV-G particles.

To generate HIV-1_NL_/SARS-CoV-2 pseudovirus, 293T cells (12 million cells/15cm plate) were co-transfected with 15.75 *µ*g CCNanoLuc/GFP, 15.75 *µ*g HIV-1_NL_ GagPol, and 5.625 *µ*g CMV-SARS-CoV-2-S, using 50ul per 15cm plate X-tremeGENE 9 (Sigma-Aldrich, 8724121001). Media was changed at 24 hr post-transfection, and supernatant was collected at 48 hr and 72 hr. Centrifuged and filtered pseudovirus was concentrated with Lenti-X-concentrator or with 40% (w/v) PEG-8000, 1.2 M NaCl, pH 7.2.

### Incucyte

Cells were imaged and analyzed using an Incucyte ZOOM (Essen BioScience). Four fields of view per well were averaged and 3-6 wells/condition were assayed in each experiment. Confluence was calculated from bright field images, GFP/Neon Green object confluence was calculated from green fluorescent images taken with 400ms exposure time, and GFP+ fractional area is the ratio of these variables.

### Luciferase assay

Following pseudovirus infection, cells were washed twice with PBS, which was subsequently aspirated. Lysis buffer (Promega, E1531) was added (50 *µ*l/well) and incubated rotating for 15 min at room temperature. NanoGlo Substrate (Promega, N1130) was diluted 1:50 in assay buffer and 25 *µ*l/well was added and incubated for an additional 15 min. Samples were transferred to a white, opaque-bottom 96W plate and luminescence was read using a BMG Labtech FLUOstar Omega plate reader.

### Clotting assays

Pooled normal human plasma was obtained from Pacific Hemostasis (VWR #95059-698). Factor X and prothrombin-deficient plasma were obtained from Haematologic Technologies (#FX-ID and #FII-ID). Russell’s Viper Venom test was performed with 10 *µ*g/ml snake venom from *Vipera russelli* (RVV, Sigma Aldrich #V2501) diluted in Tris buffer (20mM Tris HCl, 150 mM NaCl, 14 mM CaCl_2,_ pH 7.5). Pre-warmed plasma was mixed with pre-warmed dilute venom (100 *µ*l each) and monitored for visible clotting at 37 deg C. Prothrombin time was determined by mixing 100 *µ*l pre-warmed plasma with 200 *µ*l pre-warmed thromboplastin (VWR # 95059-802) and monitoring for visible clotting at 37 deg C.

---

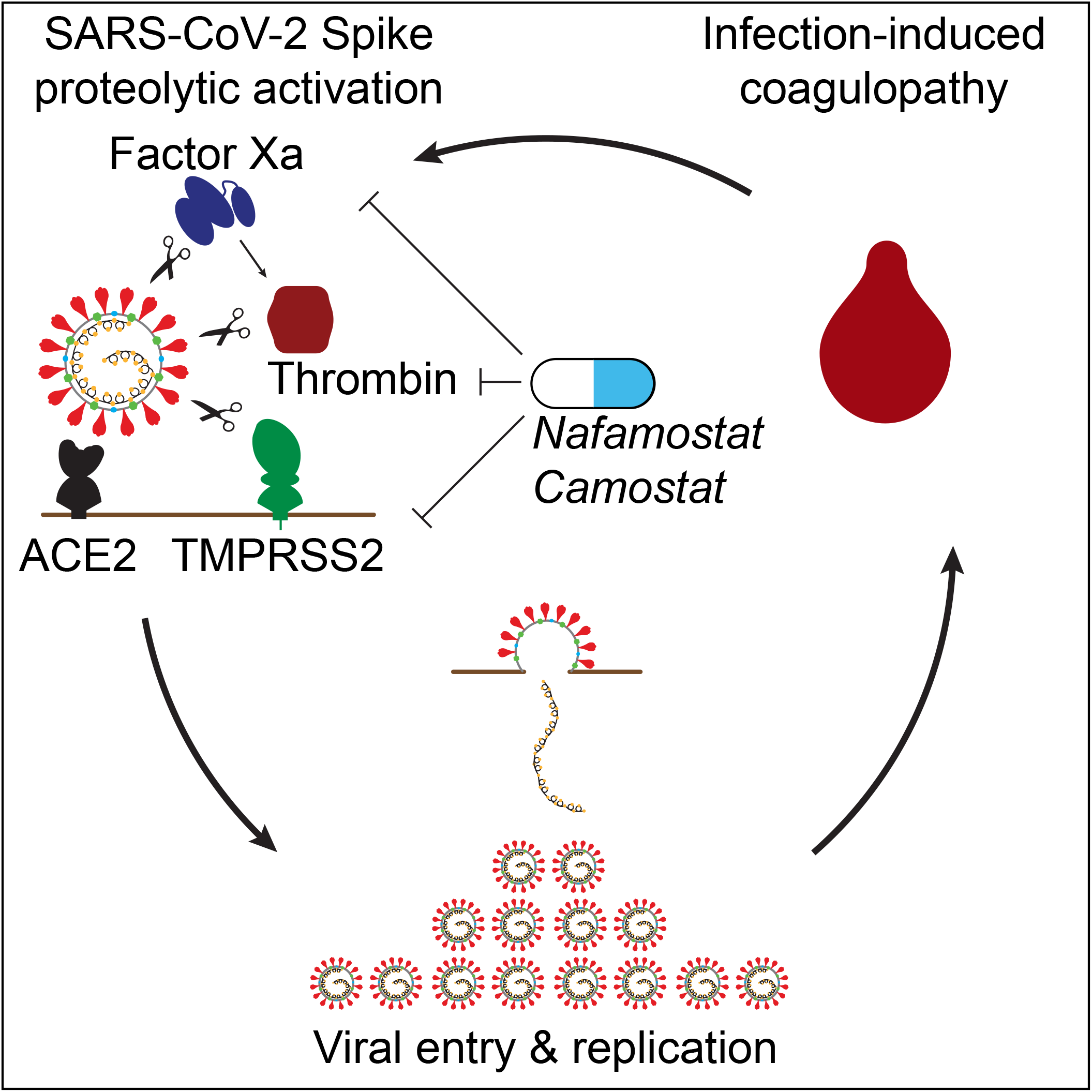
Graphical Abstract.

